# Does mechanobiology drive respiratory disease? Biomechanical induction of mucus hypersecretion in human bronchial organoids using a photocontrolled biomaterial gel

**DOI:** 10.1101/2025.06.26.661061

**Authors:** Isabel E. Uwagboe, Sharon Mumby, Iain E. Dunlop, Ian M. Adcock

## Abstract

Respiratory diseases such as COPD, IPF and severe asthma are major causes of death globally, characterized by chronic inflammation and by fibrotic biomechanical remodelling of the lung ECM. However, present treatments focus on relieving inflammation and symptoms and do not address the mechanobiological aspect. This is in great part because the role of mechanobiology in disease progression and aetiology is not well-understood, indicating a need for new investigatory models. Here we introduce a combined biomaterial and 3D-organoid model, based on a hybrid biomaterial-matrix double-network gel, whose mechanical properties are dynamically photocontrolled by the application of light. This combines basement membrane extract (Matrigel) with biocompatible polymer (poly(ethylene glycol) diacrylate), and a low-toxicity photoinitation system. We achieve rapid (<5 mins) photoinduced stiffening over the range of remodelled lung tissue (up to ∼140 kPa). Bronchosphere organoids from primary human bronchial epithelial cells, embedded within the hybrid gel, replicate airway physiology and exhibit a dynamic biological response to matrix stiffening. We show that the expression of mucus proteins Muc5AC and Muc5B is biomechanically enhanced over a period of 24 – 72 h, with in particular Muc5B showing a substantial response at 48 h after matrix stiffening. Mucus hypersecretion is a symptom of respiratory disease, and these results support the hypothesis that biomechanics is a driver of disease aetiology. We combine the photostiffened hybrid matrix gel with organoids from COPD donors, generating an advanced disease model including both cellular and biomechanical aspects. We propose this technology platform for evaluating mechanomodulatory therapeutics in respiratory disease.

## Introduction

Fibrotic lung diseases - Chronic Obstructive Pulmonary Disease (COPD), Idiopathic Pulmonary Fibrosis (IPF), and severe asthma - impose a substantial burden on society. These conditions all combine long-term chronic inflammation with remodelling of the extracellular matrix (ECM) that leads to biomechanical stiffening. COPD is the primary non-infectious cause of death globally and the second named most common lung disease in the UK[1]. In 2022, 1.4 million people were diagnosed with COPD in England with approximately 30000 people dying each year[2]. IPF has a more pronounced and rapid change in pathology; mean life expectancy from diagnosis is 3-5 years in patients aged >70 years. There are no effective treatments available and currently no cure [3, 4]. Severe asthma [defined as uncontrolled by drug treatment] is also a fibrotic disease, and its prevalence is high, often estimated at 5-10% of all asthma patients[5].

These chronic lung diseases can be treated to provide symptomatic relief: bronchodilators such as long-acting anti-muscarinic (LAMA) and long-acting beta agonists (LABA) enhance the airway patency and reduce breathlessness[4, 6] and mucolytics can aid mucus clearance from congested airways[7]. However, these drugs are palliative and do not cure the disease. A standard conceptual model of fibrotic disease induction is that inflammation onsets in response to an insult, e.g. cigarette smoke, air pollution or infection, however once the insult is removed, the inflammation does not resolve should, but rather persists into the long term. This suggests that anti-inflammatory treatment should be effective against chronic respiratory disease, and indeed topical corticosteroids are widely used and do benefit patients[8], while benralizumab has recently been successfully trialled as an anti-inflammatory treatment for COPD[9]. However while providing some relief, these anti-inflammatory treatments do not halt or reverse disease progression, showing that although fibrotic diseases originate in inflammation, other factors must also be responsible for disease continuation and progression.

This draws attention to the other key feature of fibrotic lung diseases: the remodelling of the extracellular matrix and the associated increase in biomechanical stiffness. In IPF, tissue stiffening is directly evidenced via mechanical measurements, with the Young’s modulus of IPF-patient ECM measured as ∼ 10 – 100 kPa in fibrotic regions, in comparison with 0.1 - 10 kPa for matrix from healthy donors[10]. COPD shows greater complexity within the matrix, exhibiting fibrosis and increased deposition of matrix fibres at fibrotic sites, often combined with emphysema which creates voids elsewhere[11]. Moreover, disease alters factors that impact matrix mechanical properties, including the type of matrix proteins secreted[12, 13], as well as the cross-linking of protein fibres. For example, cigarette smoking induces collagen cross-linking via advanced glycation endproducts (AGEs) and disease impacts enzymatic collagen crosslinking by lysyl oxidase [14-16].

Disease has also been shown to impact signalling pathways that are associated with mechanosensing. Specifically the Hippo/YAP-TAZ pathway shows modified activity in respiratory disease[17-19], evidenced e.g. by analysis of the most differentially expressed MiRNAs in COPD bronchoalveolar lavages[19]. Such changes in mechanosensing pathways suggest that cells are actively sensing remodelled biomechanical properties in the pathological microenvironment, and could also imply modified mechanosensing sensitivity or responsiveness by disease-phenotype cells.

These observations thus suggest the hypothesis that matrix biomechanical stiffening may be a driver of disease symptoms and progression in the airway[20]. However the evidence base for this concept consists of observational studies, and there is no direct experimental evidence that disease phenotype can be induced by mechanical effects, or of altered mechanosensing in diseased cells. To address this challenge, we here devise a new in vitro model of the airway that directly incorporates matrix stiffening. The model combines bronchosphere organoids that mimic airway biology, with a new photocontrolled biomaterial whose mechanical properties can be dynamically modified by photoillumination, enabling the sensing by airway cells of their mechanical environment to be directly measured. We use this model to investigate the biological impact of mechanical change on mucus secretion as a key symptom of respiratory disease Bronchospheres are an organoid airway model, grown in vitro using primary human epithelial cells from patients or healthy donors[21, 22]. This delivers an improved model for fibrotic disease, in comparison to organoids that are derived from induced pluripotent stem cells consist of a single or multiple layers of cells in a spherical morphology, surrounding a central [hollow] lumen[21] (Fig. 1B) that mimics the airway interior. As previously shown, bronchospheres accurately replicate the in vivo differentiation of the epithelium, with the apical layer of cells (ie the layer in contact with the lumen) including both ciliated cells and mucus-secreting goblet cells[23]. Bronchospheres are hence a good model for the cellular component of the airway.

**Figure 1.**
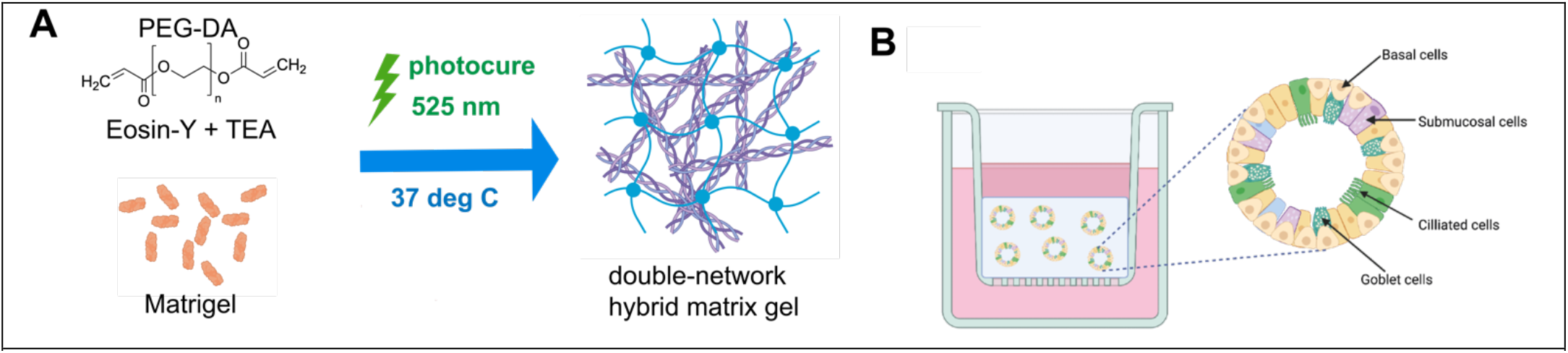
Development of a photocured double-network hybrid matrix gel. (A) Schematic of synthesis, showing formation of both synthetic network based on PEG-DA and protein-gel network based on temperature-induced Matrigel solidification. (B) Schematic showing bronchial organoids cultured in gel matrix (light blue), in transwell geometry. (Figures partially prepared using Biorender.com © biorender.com, used under licence.)

Considering the matrix aspect of the model, by far the most popular matrix-material for epithelial organoids in general – and used for bronchospheres in particular[23] - is reconstituted murine basement membrane extract from the EHS sarcoma, i.e. Matrigel or a similar commercial product (here we refer throughout to Matrigel for convenience). Matrigel is rich in laminin and provides a reasonable biochemical model for the basement membrane ECM encountered by epithelial cells in vivo[24]. However Matrigel’s reconstituted mechanical properties do not replicate those of natural ECM, and it is much less stiff than either healthy or diseased ECM[25].

Since the advent of mechanobiology, biomaterials have been used to mimic the in vitro mechanical microenvironment. 2D studies have used non-bioactive materials such as silicone elastomers or polymer hydrogels, coated with matrix proteins collagen or fibronectin[26-28], or other adhesive factors, e.g. ICAM-1[29]. In 3D organoid culture this tradition of minimally-functionalized bioinert materials is exemplified by polymer gels incorporating selected cell stimulating molecules, e.g. adhesive peptides[30], and by other systems such as synthetic peptide nanostructures[31]. Here we take a different approach, building on the rich biochemical composition of Matrigel that underpin its status as the leading gel-medium for supporting organoid culture. The stiffness of reconstituted protein gels including Matrigel can be modulated by varying the protein concentration, however this does not enable mechanobiological investigations, as mechanobiological effects cannot be distinguished from the direct impact of changing protein concentration. Hence it is desirable to develop a biomaterial that incorporates the advantages of Matrigel, but with controllable mechanics.

We here develop a hybrid gel that combines Matrigel with a synthetic polymer gel precursor. On application of light, the material stiffens, rapidly transitioning from a soft to a stiff matrix, as the synthetic polymer photocures, creating a hybrid double-network gel combining matrix and synthetic polymer (Fig. 1A). This hybrid gel exhibits the biochemistry of Matrigel, while its mechanical properties will be dominated by the synthetic polymer. The dynamic character of the measurement, whereby the organoid’s mechanical microenvironment is rapidly modified in situ, enables time-resolved measurements of the biological response, revealing time-dependent aspects of mechanobiology that would be invisible in static-gel systems, e.g. where reconstituted matrix proteins are is stiffened by addition of carbohydrates[32-34]. Using this new biomaterial system we investigate the response of human-derived bronchospheres, when the matrix stiffness is modified to values that would be associated with either healthy or fibrotic lung.

## Results

### Development of a hybrid synthetic biological matrix gel with dynamic photocontrol of stiffness

The first step was to design the double-network hybrid gel. As stated above the matrix-protein component was Matrigel, while for the synthetic network polymer we selected poly(ethylene diacrylate) (PEG-DA), building on PEG’s well-known biocompatibility as shown by e.g. its clinical application to modify drug circulation times[35]. A key challenge in developing the hybrid gel was to combine biocompatibility and lack of cytotoxic impact, with a rapid and substantial change in stiffness to span the range of healthy and diseased tissue. We selected green light (525 nm) due to anticipated lower phototoxicity than blue or UV light. As a photoinitiator system we selected eosin-Y (EY) with a co-initiator of triethanolamine (TEAO), due to its proven low phototoxicity[36]. We considered as alternatives camphorquinone, and riboflavin with TEAO, however these showed slower photocuring than EY with TEAO in preliminary tests. The concentrations of EY and TEAO were optimized for a low curing time, with the best results coming at a mass ratio of EY:TEAO = 56 (Supplementary Fig. S1) This gave a curing time of ∼ 12s using a bespoke LED setup (10W) for a 20% PEG-DA (Mw = 700 Da) gel. Incorporation of Matrigel increased this curing time presumably as the increased organic content leads to radical scavenging, hence a minimum time of 1 minute was used for the hybrid gel. As a final formulation we selected PEG-DA (Mw = 700 Da, 20% fraction), Matrigel 25%.

Photocontrol of the gel mechanical properties was demonstrated using compression testing (gel monoliths measured using 8 mm plate geometry, 50N load cell). (Fig 2B). Young’s modulus, E, as a measure of stiffness was determined by fitting the initial linear region of an engineering stress vs strain plot (Fig 2C). The stiffness of the gel, was E = 37 ± 4 kPa for 1 minute photocuring, versus E = 138 ± 23 kPa for 5 minutes photocuring (Fig 2A). This represents a very substantial increase over the value of the uncured gel, which was essentially negligible, consistent with the very low stiffness of Matrigel alone[25]. These values are within the range reported for fibrotic lung tissue[10], so that the hybrid gel constitutes an effective model of the pathological matrix microenvironment, with the curing time serving as a model parameter for the level of fibrotic stiffening.

**Figure 2.**
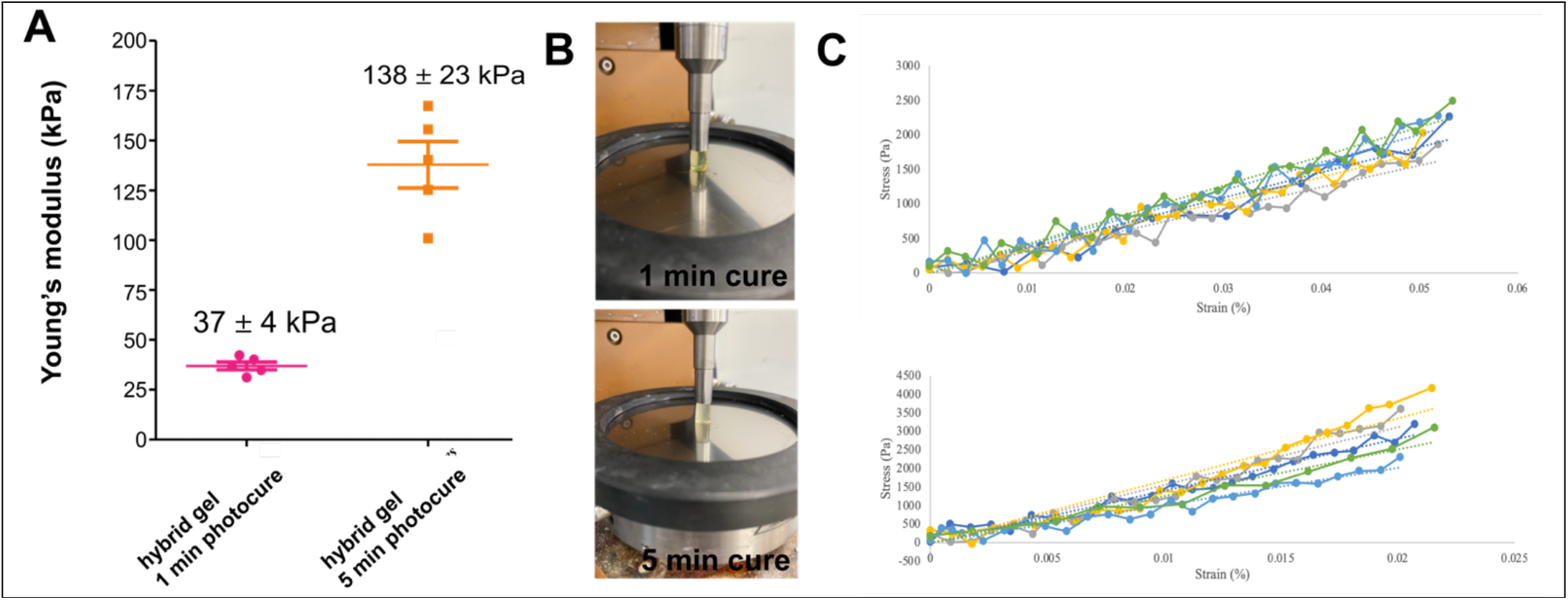
Mechanical testing of hybrid matrix gel. (A) Stiffness (Young’s modulus) of gel with different photocuring times measured using (B) compression testing. (C) Confirmation of linearity in engineering stress-strain compression curves used to derive Young’s modulus: (top panel) 1 min photocure; (bottom panel) 5 mins photocure (symbols show data from individual samples and dotted lines fits.)

### Human bronchial organoids are viable and maintain structure in photostiffened hybrid matrix gel

To establish the photocured hybrid gel matrix as a model microenvironment for biomechanical investigations of the respiratory system, bronchosphere organoids (established in Matrigel) were transferred into the hybrid gel precursor solution, in a transwell geometry (Fig. 1B), which was then photocured. Confocal fluorescence microscopy was used to investigate organoid viability, based on the maintenance of organoid structure, nuclear integrity, and the presence of proliferating cells. The structural integrity of organoids was shown to be maintained by nuclear imaging over 72h, with no sign of disruption by any toxic effect (Fig 3).

**Figure 3.**
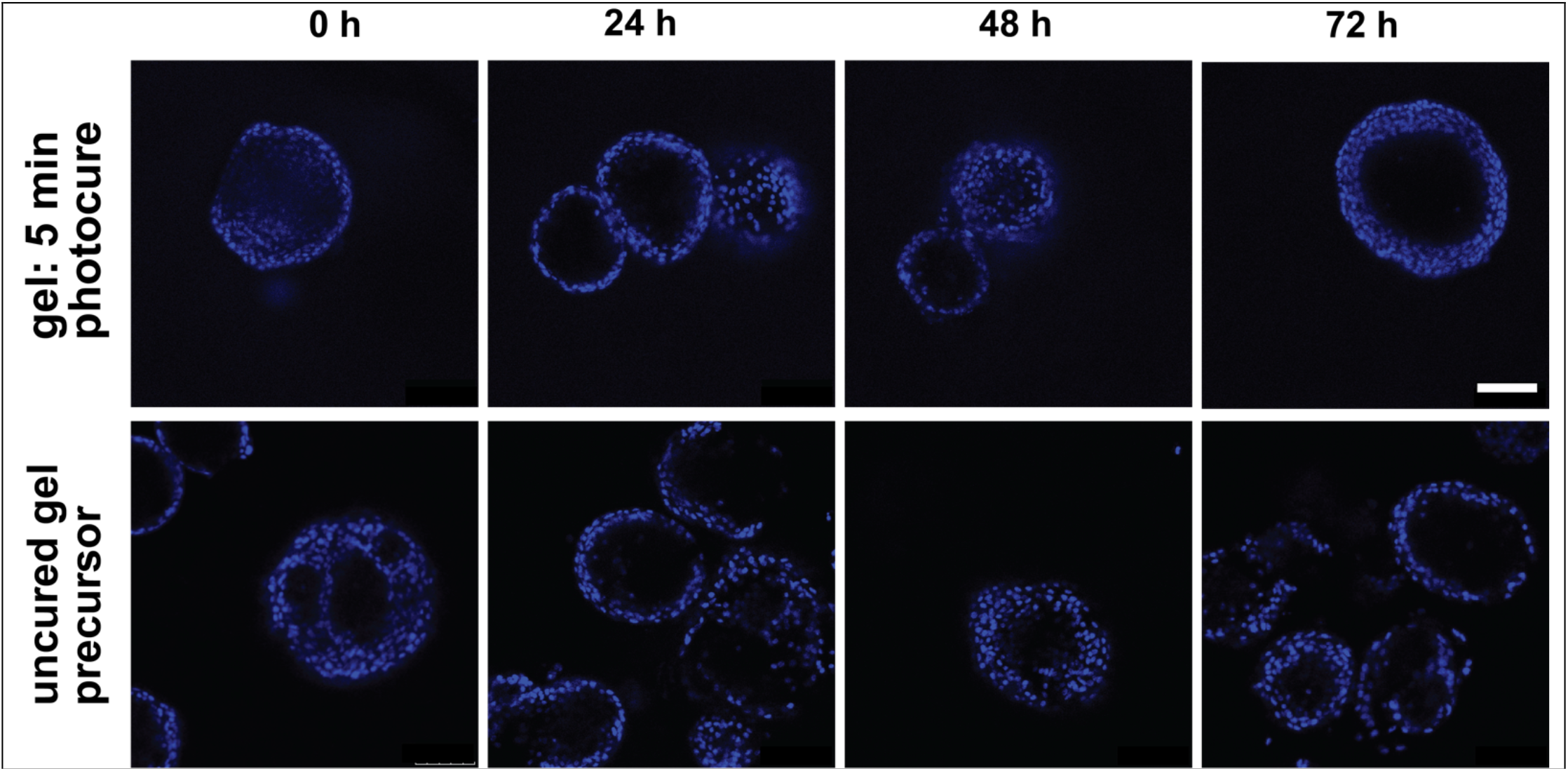
Culture of bronchial organoids in photostiffened hybrid matrix gel. Time course showing integrity of cell nuclei and organoid structure from 0 – 72 hours in gel (5 minute photocured) and uncured gel precursor (Blue = DAPI). Scale bar 100μm. (Images from distinct experiments for each time point.)

### Mucus secretion in human bronchial organoids is biomechanically modulated by matrix stiffening

We next investigated the impact of matrix-stiffening on mucus secretion. The secretion of mucus proteins MUC5AC and MUC5B by bronchosphere in stiffened hybrid gels vs control was investigated by confocal immunofluorescence microscopy (fixed and permeabilized samples, Fig. 4). Both of these mucus proteins are secreted in organoids, and this is observed across all matrix types: Matrigel and the 1 min and 5 min photocured hybrid gels. The secretion of mucus shows that these organoids contain epithelial cells of the goblet sub-type, indicating further they are a good representation of the in vivo differentiated epithelium (Fig 1B). It is interesting to note that there is no clear spatial relationship between secretion of the two mucus proteins: in some cases they are secreted in the same location, and in some cases in separate locations. This is consistent with the results of a prior investigation on an air-liquid interface model of bronchial epithelium, which showed the existence of cells that secrete only one or the other of these proteins, as well as some cells that secrete both (data from Human Lung Atlas core dataset[37]; Suppl. Fig. S2). A 3D reconstructed image showing mucus secretion in an organoid in a hybrid matrix gel is appended (Suppl. Video S1).

**Figure 4.**
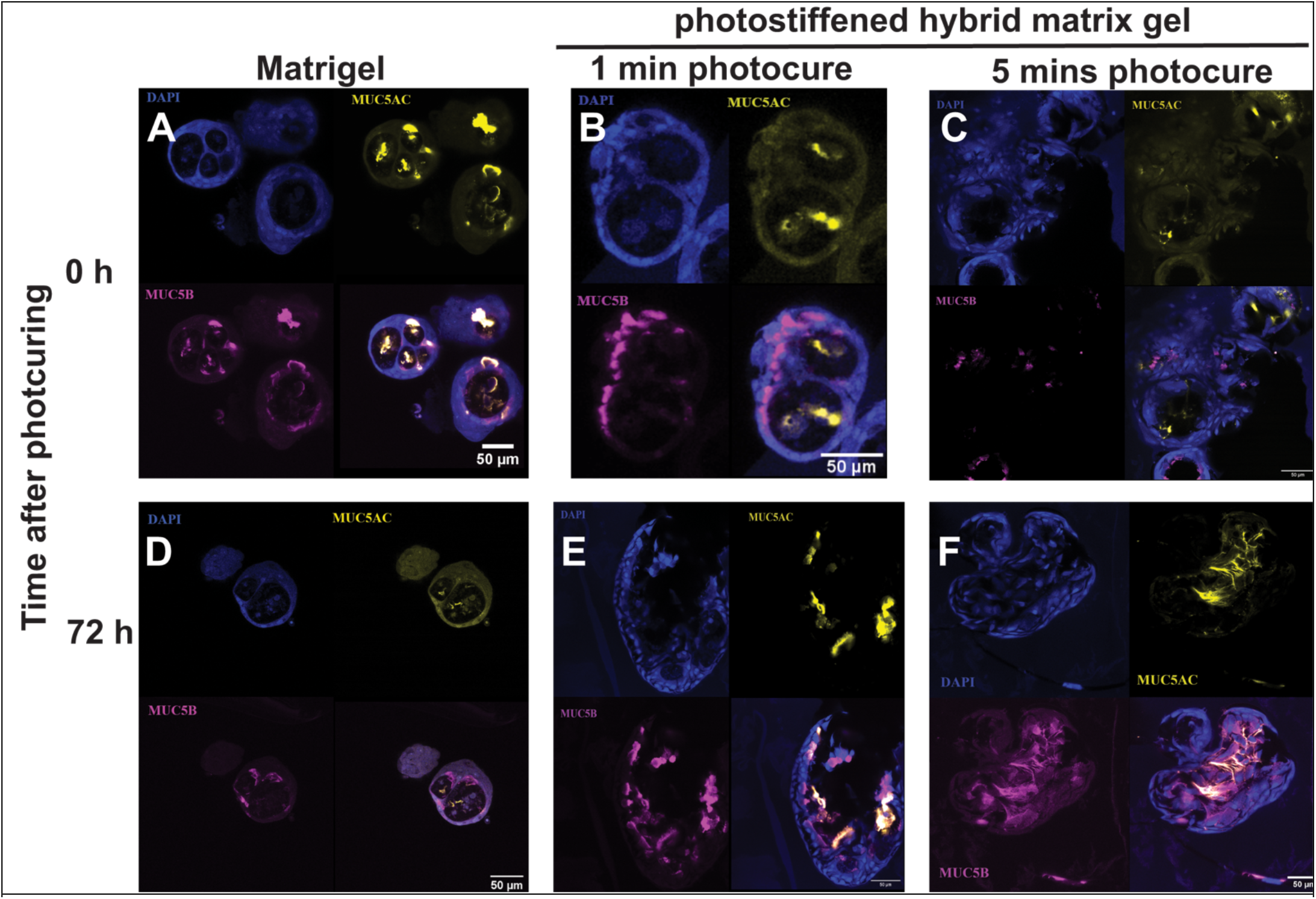
Mucus secretion in human bronchial organoids after photosti8ening of matrix. Confocal immunofluorescence imaging of MUC5AC (Yellow) and MUC5B (pink) along with cell nuclei (DAPI, blue). Images are shown in Matrigel as a control (A, D), photostiffened hybrid matrix gel after 1 minute photocuring (B,E) and 5 minutes photocure, at time points of 0 h (A,B,C) and 72 h after photocuring/light exposure (D, E, F).

Quantitative evaluation of mucus secretion in photostiffened gels versus control was performed using quantitative PCR (q-PCR, determining the amount of messenger RNA) to evaluate expression of the relevant genes (Fig. 5). Expression was determined dynamically as a function of time after photocuring: an important benefit of the near-instant photocuring proceses (in comparison to culture times). The overall value for each condition was normalized to control organoids in uncured gel precursor, so that the graph effectively shows fold-change for the photostiffened gels versus uncured (after initial normalization to an average of housekeeping genes).

**Figure 5.**
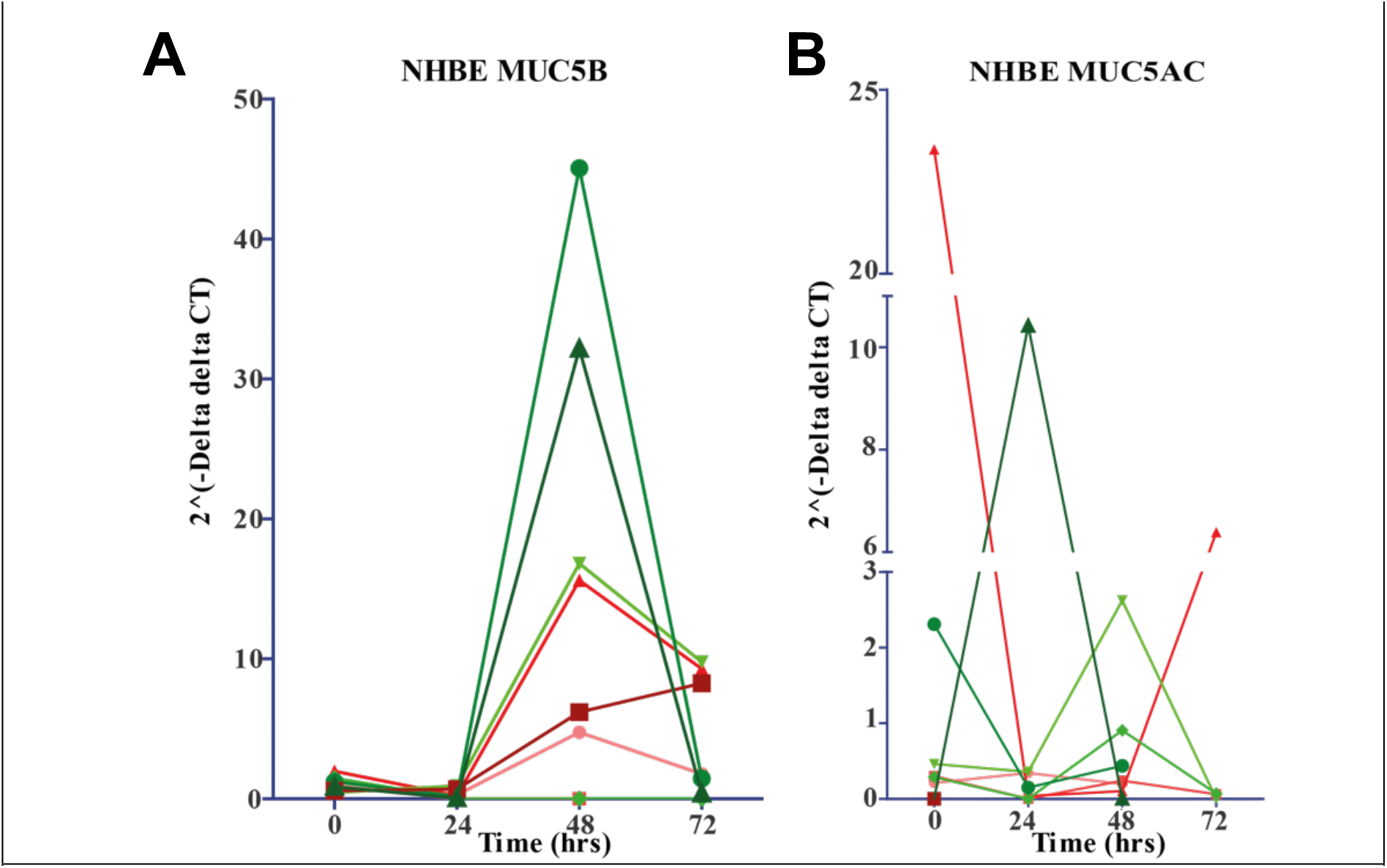
Biomechanical modulation of mucus protein expression in human bronchial organoids. Plots show expression analysed by q-PCR of MUC5B (A) and MUC5AC (B), for organoids in gels photocured for 1 minute (red) and 5 minutes (green), with symbol indicating organoids from distinct human donors (n=4), as a function of incubation time after photocuring (0-72h). All values are normalized to a control sample which was cultured in the hybrid gel precursor but without photocuring.

The impact of stiffening the matrix is to increase mucus secretion. For MUC5B, the effect is consistent and sensitive at 48h after stiffening, with the stiffest (5 min photocure) gels showing high levels of increase in MUC5B secretion (3/4 donors), while intermediate stiffness (1 min photocure) gels show a lesser but still substantial effect (Fig 5A). For MUC5AC, secretion increases in the stiffest gel (5 min cure) but not the intermediate stiffness (1 min cure, ignoring an outlier with anomalously high initial stiffness) (Fig. 5B). Here, there is some variation in the timepoint of maximum stiffening. Indeed, across the two proteins, we see dynamic variation in mucus secretion, in particular for MUC5B we see essentially no change after 24h, but the increase in mucus secretion reaches a high maximum level at 48h.

### Photostiffened hybrid gel can be used as a scaffold for organoid investigations of COPD

Finally, we explored whether the photostiffened gel can be used to generate an advanced in vitro 3D model of a respiratory disease. A realistic model of respiratory disease in vitro should combine disease-altered cells, together with a matrix that represents the remodelled/stiffened matrix associated with disease. As a proof of concept, organoids were created using the same protocols as before, but using primary human epithelial cells derived from donors with COPD, and were embedded in the photostiffened hybrid gel. Confocal microscopy was used to image cell nuclei and mucus secretion for gels of multiple stiffnesses at 0 h and 48 h after photocuring Fig. 6). These images show the maintenance of organoid integrity and mucus secretion, as for healthy organoids. Hence these results indicate that the generation of a mechanically and cellularly realistic 3D in vitro model of COPD.

**Figure 6.**
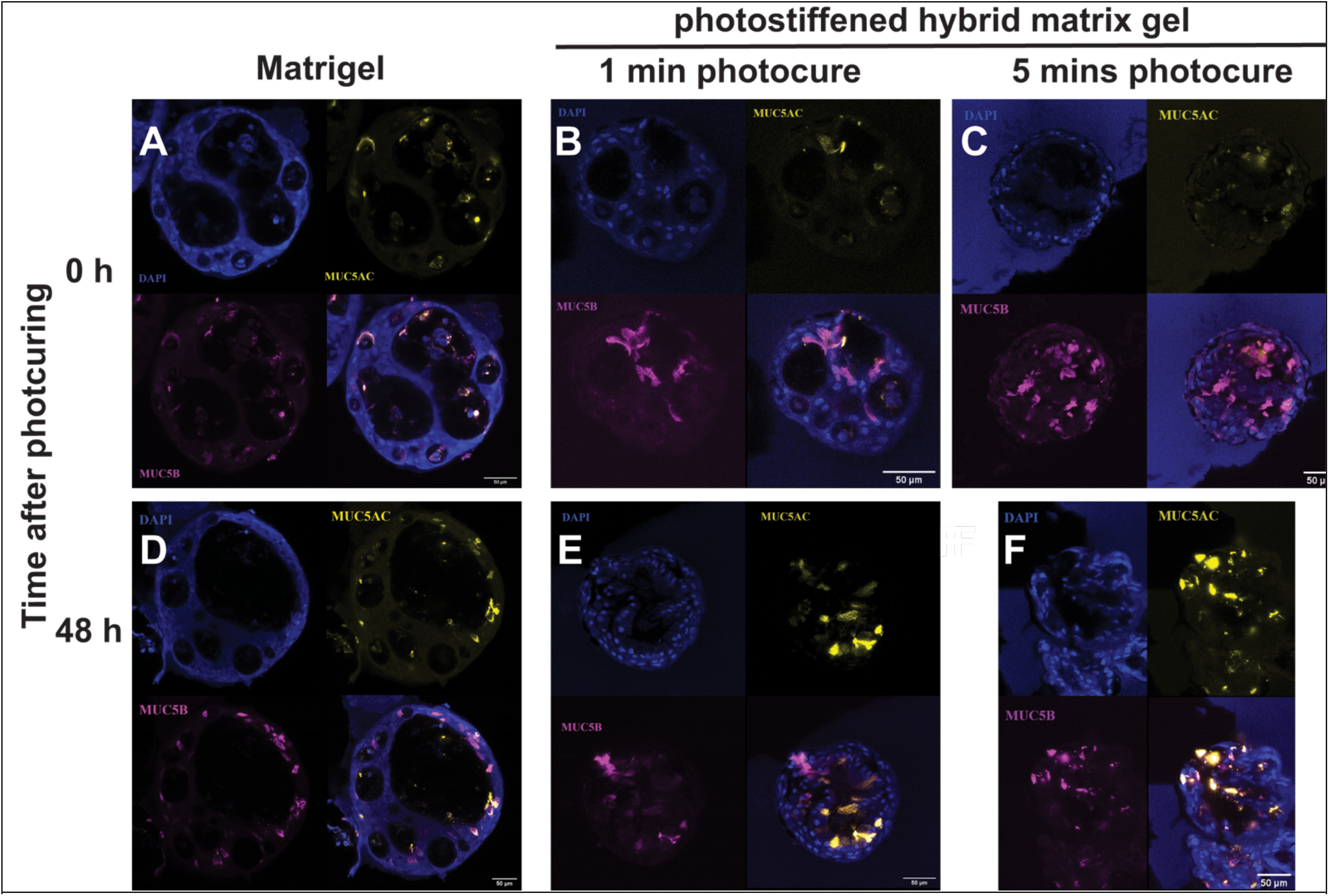
Culture of COPD-derived bronchial epithelial organoids in photostiffened hybrid matrix gel. Organoids grown from primary human bronchial epithelial cells (CHBEs), and embedded into photostiffened hybrid matrix gel in the same way as for normal organoids above. Confocal immunofluorescence imaging of MUC5AC (Yellow) and MUC5B (pink) along with cell nuclei (DAPI, blue). Images are shown in Matrigel as a control (A, D), photostiffened hybrid matrix gel after 1 minute photocuring (B,E) and 5 minutes photocure, at time points of 0 h (A,B,C) and 48 h after photocuring/light exposure (D, E, F).

## Discussion

Our results establishes experimentally for the first time that mucus hypersecretion, a key disease symptom in both COPD and IPF, can be directly induced by mechanical change in the matrix mechanical properties. This suggests that stiffening of the extracellular matrix may be a driver of fibrotic lung disease, rather than a purely passive output of the disease process. This gives support to a recently proposed conceptual model of fibrosis [20, 38]. In this view, the initial insult and inflammation lead to changes in matrix mechanical properties, in particular increasing stiffness. These altered mechanical properties are then sensed by epithelial cells, which respond by generating a disease phenotype, including potentially more inflammation, leading to disease progression. Hence even if anti-inflammatory treatment reduces inflammation, the altered mechanical properties remain, and will play a role in driving disease progression, and in restarting it if treatment is stopped. While preliminary, our results give clear support to this model. The light-induced nature of the stiffening, permitting both dynamic variation of matrix stiffness, and the use of a comparable uncured control, makes the mechanically-controlled nature of this change very clear. The technology we have created can thus function as an effective platform for mechanobiological investigation of organoid systems, in respiratory disease and beyond.

Furthermore, the construction of an ex vivo organoid system with the ability to biomechanically induce a disease symptom – mucus hypersecretion – opens the possibility to test drugs that inhibit or modulate mechanosensing, with the aim of relieving this symptom in patients. This is significant because mucus hypersecretion is a key symptom of both COPD and IPF, and responsible for many of the adverse effects experienced by patients with these conditions [39]. Current treatments address the effects but not the underlying biological causes of mucus secretion. Hence the concept of mechanosensing modulation as a potential mechanism of action is of value, especially since reagents that modulate mechanosensing signalling pathways are increasingly in pre-clinical investigation across a range of contexts. Furthermore we have shown that the photostiffened matrix gel can be combined with COPD organoids to generate a combined mechanical and biological in vitro disease model.

From a biophysics perspective, our results demonstrate a new instance of biomechanical response in epithelial tissue. Across many tissues, it has now been shown that cells sense forces and the mechanical properties of their environment, and these impact complex cellular behaviours including stem cell differentiation[40, 41], cell migration[42] and metastasis[43], and immune cell activation[29]. An emerging challenge in the field is to understand how such mechanosensing impacts disease, and our results make a contribution to this. Technologically, we have introduced a new and valuable mechanically-controlled biomaterial to the mechanobiology toolbox. As mentioned above there is a need for technologies that modulate matrix stiffness across a wide range of values without changing the concentration of matrix biomolecules, and our photostiffened gel achieves this. Alternative biomaterial approaches that have been reported include mixing matrix extract with carbohydrates such as agarose and alginate[32-34], however in this case the stiffness is determined from the start, and cannot be altered dynamically as in our photostiffened material. The PEG-DA-based approach is exceptionally straightforward to implement, and does not require prior chemical modification of the matrix proteins[44]. Of equal importance, we have used as a protein source Matrigel, which is readily available commercially in contrast to patient-derived matrices[44] and which leads to a laminin-rich biomaterials suited to epithelial cells, rather than a pure collagen gel[45, 46] which does not effectively represent the basement membrane.

In conclusion, this study shows for the first time the induction of a key respiratory disease secretion - mucus hypersecretion – by mechanically stiffening the matrix. This is achieved by introducing a new bioengineered in vitro model for lung disease that models pathological matrix stiffening along with the biology of the epithelium. These results help shed light on the biology of respiratory disease, and open the path towards new mechanomodulatory therapies.

## Methods

### Cells and organoid generation

Normal human primary bronchial epithelial cells (NHBEs) were sourced from Lonza Cell stocks (*Lonza, Basel, Switzerland*) and Promocell (*Promocell, Heidelberg, Germany*) or from our internal cell bank (*National Heart and Lung Institute, Imperial College London, London, UK*). Donors were 50% Male 50% female and in the age range 38 – 69 years (mean 55 years) (donor details see Suppl. Table S1 Human primary bronchial epithelial cells from COPD donors CHBEs) were similarly sourced (Suppl. Table S1).

A co-culture system was initially used to create the lung organoids, using NIH/3T3 mouse fibroblasts and human bronchial epithelial cells as previously described[32, 47, 48]. Feeder layers were seeded, cultured to ∼70% confluency and then growth arrested in order to introduce the human bronchial epithelial cells. Once passaged, the bronchial epithelial cells were seeded as single cells on a bed of 25% Matrigel (Corning Life Sciences, Amsterdam, Netherlands) and fed on day 3, 8, 14 and then weekly until mature bronchospheres were formed. Bronchospheres were harvested on day 24-27.

### Organoid implantation into biomaterial hybrid gel

Gel precursor solution was created from polyethylene diacrylate (PEG-DA, Mw 700 Da, Sigma-Aldrich/Merck, Dorset, U.K, 20 vol% final concentration in precursor solution); culture media (composition see Suppl. Table S2; 35 vol% final concentration); Matrigel (Corning Life Sciences B.V., Amsterdam, The Netherlands; 25 vol%), photoinitiator system: triethanolamine solution (0.5% v/v in sterile water, 10% by volume) and Eosin Y solution (0.01% w/v in ethanol; 10 vol%) both from Sigma-Aldrich. Reagents were added in the following order with pipette mixing: culture media containing organoids; triethanolamine; Eosin-Y; Matrigel (stored on. ice to keep it liquid). The suspension was transferred to a 24-well transwell insert (Sarstedt AG & Co. KG, Germany) which was then placed in a 12-well plate to ensure plastic-plastic contact at the bottom for improved light transmission, and sealed with parafilm. Light curing used a 10W Pure Green light wavelength (525nm) LED array, with the microwell placed directly on top of the light source. Photocuring time was either 1 min or 5 min depending on desired gel stiffness, and also uncured controls were used (no light exposure). After curing, the transwell insert was placed in a 24-well plate, and culture media was added both above the gel and to the lower part of the transwell assembly. Some organoid samples were harvested immediately after curing (0 hours), otherwise, organoids were then cultured for either 24, 48 or 72 hours.

### Mechanical characterization of biomaterial hybrid gel

Mechanical characterization used monolith samples of photostiffened hybrid matrix gel prepared without organoids. These were subject to compression tests (Discovery Hybrid rheometer, TA Instruments, Delaware, USA, 8mm diameter parallel plate geometry, axial force mode, 50N effective load cell). Young’s modulus was determined by fitting the initial linear region of an engineering stress vs strain curve (Fig 2C).

### Confocal Imaging

Confocal images were taken by using whole mount sample staining as used previously[49]. Samples were fixed, permeabilised and stained for cell subtypes using cell-specific antibodies (Suppl. Table S3). Tissue clearing was used to clarify images generated.

### Quantitative PCR (q-PCR) measures of gene expression

Q-PCR was used to quantify the expression of selected genes according to previously used approaches[50], normalization as per main text.

## Acknowledgements

This work was funded by an EPSRC Doctoral Partnership studentship to IEU (EP/N509486/1), and the use of equipment funded by EP/M000044/1. We acknowledge support from the South Kensington Flow Cytometry Facility, and the Facility for Imaging with Light Microscopy, both at Imperial College London. We thank Mark Oxborrow for assistance with setting up the photocuring apparatus.

## Data Availability Statement

Data has been included where possible in the article or supplementary information; otherwise it is available from the corresponding authors on request.

## References

[1] Asthma+LungUK, COPD in the UK: Delayed diagnosis and unequal care. Executive summary and recommendations, 2022.

[2] A. Kulakiewicz, M. Macdonald, C. Baker, Support for people with chronic obstructive pulmonary disease., House of Commons Library, UK Parliament, United Kingdom., 2021.

[3] T.M. Maher, E. Bendstrup, L. Dron, J. Langley, G. Smith, J.M. Khalid, H. Patel, M. Kreuter, Global incidence and prevalence of idiopathic pulmonary fibrosis, Respir. Res. 22(1) (2021) 197.

[4] L.G. Spencer, M. Loughenbury, N. Chaudhuri, M. Spiteri, H. Parfrey, Idiopathic pulmonary fibrosis in the UK: analysis of the British Thoracic Society electronic registry between 2013 and 2019, ERJ Open Res. 7(1) (2021).

[5] K.F. Chung, S.E. Wenzel, J.L. Brozek, A. Bush, M. Castro, P.J. Sterk, I.M. Adcock, E.D. Bateman, E.H. Bel, E.R. Bleecker, L.P. Boulet, C. Brightling, P. Chanez, S.E. Dahlen, R. Djukanovic, U. Frey, M. Gaga, P. Gibson, Q. Hamid, N.N. Jajour, T. Mauad, R.L. Sorkness, W.G. Teague, International ERS/ATS guidelines on definition, evaluation and treatment of severe asthma, Eur. Respir. J. 43(2) (2014) 343–73.

[6] I. Uwagboe, I.M. Adcock, F. Lo Bello, G. Caramori, S. Mumby, New drugs under development for COPD, Minerva Med. 113(3) (2022) 471–496.

[7] P. Poole, K. Sathananthan, R. Fortescue, Mucolytic agents versus placebo for chronic bronchitis or chronic obstructive pulmonary disease, Cochrane Database of Systematic Reviews (5) (2019).

[8] I.M. Adcock, G. Caramori, P.A. Kirkham, Strategies for improving the efficacy and therapeutic ratio of glucocorticoids, Cur Opin. Pharmacol. 12(3) (2012) 246–251.

[9] S. Ramakrishnan, R.E.K. Russell, H.R. Mahmood, K. Krassowska, J. Melhorn, C. Mwasuku, I.D. Pavord, L. Bermejo-Sanchez, I. Howell, M. Mahdi, S. Peterson, T. Bengtsson, M. Bafadhel, Treating eosinophilic exacerbations of asthma and COPD with benralizumab (ABRA): a double-blind, double-dummy, active placebo-controlled randomised trial, Lancet Respir. Med. (2024).

[10] A.J. Booth, R. Hadley, A.M. Cornett, A.A. Dreffs, S.A. Matthes, J.L. Tsui, K. Weiss, J.C. Horowitz, V.F. Fiore, T.H. Barker, B.B. Moore, F.J. Martinez, L.E. Niklason, E.S. White, Acellular normal and fibrotic human lung matrices as a culture system for in vitro investigation, Am J Respir Crit Care Med 186(9) (2012) 866–76.

[11] M. Karakioulaki, E. Papakonstantinou, D. Stolz, Extracellular matrix remodelling in COPD, Eur Respir Rev 29(158) (2020).

[12] G. Burgstaller, B. Oehrle, M. Gerckens, E.S. White, H.B. Schiller, O. Eickelberg, The instructive extracellular matrix of the lung: basic composition and alterations in chronic lung disease, Eur Respir J 50(1) (2017).

[13] M.L. Decaris, M. Gatmaitan, S. FlorCruz, F. Luo, K. Li, W.E. Holmes, M.K. Hellerstein, S.M. Turner, C.L. Emson, Proteomic Analysis of Altered Extracellular Matrix Turnover in Bleomycin-induced Pulmonary Fibrosis, Mol Cell Proteomics 13(7) (2014) 1741–1752.

[14] C. Onursal, E. Dick, I. Angelidis, H.B. Schiller, C.A. Staab-Weijnitz, Collagen Biosynthesis, Processing, and Maturation in Lung Ageing, Front Med-Lausanne 8 (2021).

[15] G. Tjin, E.S. White, A. Faiz, D. Sicard, D.J. Tschumperlin, A. Mahar, E.P.W. Kable, J.K. Burgess, Lysyl oxidases regulate fibrillar collagen remodelling in idiopathic pulmonary fibrosis, Dis Model Mech 10(11) (2017) 1301–1312.

[16] V. Aumiller, B. Strobel, M. Romeike, M. Schuler, B.E. Stierstorfer, S. Kreuz, Comparative analysis of lysyl oxidase (like) family members in pulmonary fibrosis, Sci Rep 7(1) (2017) 149.

[17] P. Kachroo, J.D. Morrow, A.T. Kho, C.A. Vyhlidal, E.K. Silverman, S.T. Weiss, K.G. Tantisira, D.L. DeMeo, Co-methylation analysis in lung tissue identifies pathways for fetal origins of COPD, Eur Respir J 56(4) (2020).

[18] H.J. Xie, L.Q. Wu, Z.A. Deng, Y.T. Huo, Y.X. Cheng, Emerging roles of YAP/TAZ in lung physiology and diseases, Life Sci 214 (2018) 176–183.

[19] N. Cerón-Pisa, A. Iglesias, H. Shafiek, A. Martín-Medina, M. Esteva-Socias, J. Muncunill, A. Fleischer, J. Verdú, B.G. Cosío, J. Sauleda, Hsa-Mir-320c, Hsa-Mir-200c-3p, and Hsa-Mir-449c-5p as Potential Specific miRNA Biomarkers of COPD: A Pilot Study, Pathophysiology-Base 29(2) (2022) 143–156.

[20] M.A.T. Freeberg, A. Perelas, J.K. Rebman, R.P. Phipps, T.H. Thatcher, P.J. Sime, Mechanical Feed-Forward Loops Contribute to Idiopathic Pulmonary Fibrosis, Amer. J. Pathol. 191(1) (2021) 18–25.

[21] P. Beri, Y.J. Woo, K. Schierenbeck, K. Chen, S.W. Barnes, O. Ross, D. Krutil, D. Quackenbush, B. Fang, J. Walker, W. Barnes, E.Q. Toyama, A high-throughput cigarette smoke-treated bronchosphere model for disease-relevant phenotypic compound screening, PLoS One 18(6) (2023) e0287809.

[22] J. van der Vaart, H. Clevers, Airway organoids as models of human disease, J Intern Med 289(5) (2021) 604–613.

[23] Y.C. Zheng, Q.Y. Tian, H.W. Yang, Y.D. Cai, J.X. Zhang, Y.F. Wu, S. Zhu, Z.C. Qiu, Y.M. Lin, J.Q. Hong, Y. Zhang, D. Dockrell, S.H. Ma, Identification of Nicotinic Acetylcholine Receptor for N-Acetylcysteine to Rescue Nicotine-induced Injury Using Beating Cilia in Primary Tissue Derived Airway Organoids, Adv Sci (2024).

[24] H.K. Kleinman, M.L. Mcgarvey, J.R. Hassell, V.L. Star, F.B. Cannon, G.W. Laurie, G.R. Martin, Basement-Membrane Complexes with Biological-Activity, Biochemistry-Us 25(2) (1986) 312–318.

[25] K. Slater, J. Partridge, H. Nandivada, Tuning the Elastic Moduli of Corning® Matrigel® and Collagen I 3D Matrices by Varying the Protein Concentration, Corning, Corning Application Note.

[26] J.L. Young, A.J. Engler, Hydrogels with time-dependent material properties enhance cardiomyocyte differentiation in vitro, Biomaterials 32(4) (2011) 1002–9.

[27] B. Trappmann, J.E. Gautrot, J.T. Connelly, D.G.T. Strange, Y. Li, M.L. Oyen, M.A.C. Stuart, H. Boehm, B.J. Li, V. Vogel, J.P. Spatz, F.M. Watt, W.T.S. Huck, Extracellular-matrix tethering regulates stem-cell fate, Nat Mater 11(7) (2012) 642–649.

[28] G.K. Toworfe, R.J. Composto, C.S. Adams, I.M. Shapiro, P. Ducheyne, Fibronectin adsorption on surface-activated poly(dimethylsiloxane) and its effect on cellular function, J Biomed Mater Res A 71a(3) (2004) 449–461.

[29] S. Sasidharan, D.M. Davis, I.E. Dunlop, Bioinspired Materials for Immunoengineering of T Cells and Natural Killer Cells, Adv Funct Mater 34(35) (2024).

[30] L.E.E. Jansen, H. Kim, C.L. Hall, T.P. McCarthy, M.J. Lee, S.R. Peyton, A poly(ethylene glycol) three-dimensional bone marrow hydrogel, Biomaterials 280 (2022).

[31] L.A. Castillo-Diaz, J.E. Gough, A.F. Miller, A. Saiani, RGD-functionalised self-assembling peptide hydrogel induces a proliferative profile in human osteoblasts in vitro, J Pept Sci (2024) e3653.

[32] T.G. Güney, A.M. Herranz, S. Mumby, I.E. Dunlop, I.M. Adcock, Epithelial-stromal cell interactions and extracellular matrix mechanics drive the formation of airway-mimetic tubular morphology in lung organoids, Iscience 24(9) (2021).

[33] S.R. Moxon, N.J. Corbett, K. Fisher, G. Potjewyd, M. Domingos, N.M. Hooper, Blended alginate/collagen hydrogels promote neurogenesis and neuronal maturation, Mater Sci Eng C Mater Biol Appl 104 (2019) 109904.

[34] S. Deng, Y. Zhu, X. Zhao, J. Chen, R.S. Tuan, H.F. Chan, Efficient fabrication of monodisperse hepatocyte spheroids and encapsulation in hybrid hydrogel with controllable extracellular matrix effect, Biofabrication 14(1) (2021).

[35] M. Hamidi, A. Azadi, P. Rafiei, Pharmacokinetic consequences of pegylation, Drug Deliv 13(6) (2006) 399–409.

[36] A. Fu, K. Gwon, M. Kim, G. Tae, J.A. Kornfield, Visible-light-initiated thiol-acrylate photopolymerization of heparin-based hydrogels, Biomacromolecules 16(2) (2015) 497–506.

[37] H.B. Schiller, D.T. Montoro, L.M. Simon, E.L. Rawlins, K.B. Meyer, M. Strunz, F.A. Vieira Braga, W. Timens, G.H. Koppelman, G.R.S. Budinger, J.K. Burgess, A. Waghray, M. van den Berge, F.J. Theis, A. Regev, N. Kaminski, J. Rajagopal, S.A. Teichmann, A.V. Misharin, M.C. Nawijn, The Human Lung Cell Atlas: A High-Resolution Reference Map of the Human Lung in Health and Disease, Am J Respir Cell Mol Biol 61(1) (2019) 31–41.

[38] Y. Long, Y.D. Niu, K.N. Liang, Y.N. Du, Mechanical communication in fibrosis progression, Trends Cell Biol. 32(1) (2022) 70–90.

[39] R. Balsamo, L. Lanata, C.G. Egan, Mucoactive drugs, Eur Respir Rev 19(116) (2010) 127–33.

[40] K.H. Vining, D.J. Mooney, Mechanical forces direct stem cell behaviour in development and regeneration, Nat Rev Mol Cell Bio 18(12) (2017) 728–742.

[41] H. Atcha, Y.S. Choi, O. Chaudhuri, A.J. Engler, Getting physical: Material mechanics is an intrinsic cell cue, Cell Stem Cell 30(6) (2023) 750–765.

[42] M. Mathieu, A. Isomursu, J. Ivaska, Positive and negative durotaxis -mechanisms and emerging concepts, J Cell Sci 137(8) (2024).

[43] E. Cambria, M.F. Coughlin, M.A. Floryan, G.S. Offeddu, S.E. Shelton, R.D. Kamm, Linking cell mechanical memory and cancer metastasis, Nat Rev Cancer 24(3) (2024) 216–228.

[44] M. Nizamoglu, R.H.J. de Hilster, F. Zhao, P.K. Sharma, T. Borghuis, M.C. Harmsen, J.K. Burgess, An in vitro model of fibrosis using crosslinked native extracellular matrix-derived hydrogels to modulate biomechanics without changing composition, Acta Biomater 147 (2022) 50–62.

[45] Z. Bao, M. Gao, X. Fan, Y. Cui, J. Yang, X. Peng, M. Xian, Y. Sun, R. Nian, Development and characterization of a photo-cross-linked functionalized type-I collagen (Oreochromis niloticus) and polyethylene glycol diacrylate hydrogel, Int J Biol Macromol 155 (2020) 163–173.

[46] L. Tytgat, A. Dobos, M. Markovic, L. Van Damme, J. Van Hoorick, F. Bray, H. Thienpont, H. Ottevaere, P. Dubruel, A. Ovsianikov, S. Van Vlierberghe, High-Resolution 3D Bioprinting of Photo-Cross-linkable Recombinant Collagen to Serve Tissue Engineering Applications, Biomacromolecules 21(10) (2020) 3997–4007.

[47] C.R. Butler, R.E. Hynds, K.H.C. Gowers, D.D.H. Lee, J.M. Brown, C. Crowley, V.H. Teixeira, C.M. Smith, L. Urbani, N.J. Hamilton, R.M. Thakrar, H.L. Booth, M.A. Birchall, P. De Coppi, A. Giangreco, C. O’Callaghan, S.M. Janes, Rapid Expansion of Human Epithelial Stem Cells Suitable for Airway Tissue Engineering, Am J Resp Crit Care 194(2) (2016) 156–168.

[48] R.E. Hynds, A. Ben Aissa, K.H.C. Gowers, T.B.K. Watkins, L. Bosshard-Carter, A.J. Rowan, S. Veeriah, G.A. Wilson, S.A. Quezada, C. Swanton, S.M. Janes, T. Consortium, Expansion of airway basal epithelial cells from primary human non-small cell lung cancer tumors, Int J Cancer 143(1) (2018) 160–166.

[49] M.Z. Nikolic, E.L. Rawlins, Lung Organoids and Their Use To Study Cell-Cell Interaction, Curr Pathobiol Rep 5(2) (2017) 223–231.

[50] S. Mumby, F. Perros, J. Grynblat, G. Manaud, A. Papi, P. Casolari, G. Caramori, M. Humbert, S. John Wort, I.M. Adcock, Differential responses of pulmonary vascular cells from PAH patients and controls to TNFalpha and the effect of the BET inhibitor JQ1, Respir Res 24(1) (2023) 193.

