## Supplementary information for "Does mechanobiology drive respiratory disease? Biomechanical induction of mucus hypersecretion in human bronchial organoids using a photocontrolled biomaterial gel"

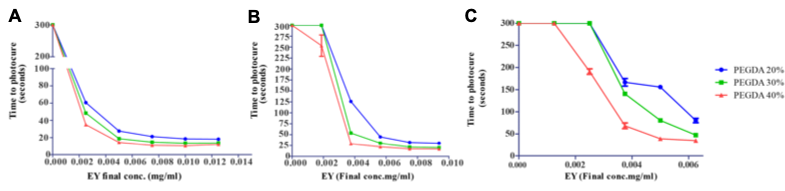

***Supplementary Figure S1. Optimization of photoinitiator system.*** *Time taken for PEG-DA 700MW hydrogel scaffold to fully photocure using photoinitiator eosin Y (EY) and co-initiator triethanolamine (TEAO). Gels were photocured for >300s. Where ‘300s’ is plotted on graph, no curing has taken place. (A and D) EY:TEAO, at the ratio of 1:5.62. (B and E) EY:TEAO, at the ratio of 1:37.47. (C and F) EY:TEAO, at the ratio of 1:22.48 (Mean and standard error in mean, n=3 repeats).*

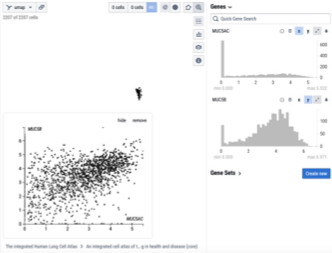

**Supplementary Figure S2 Co-expression of MUC5AC and MUC5B.** Display of data from Human Lung Atlas core dataset; originally generated by single-cell RNA-sequencing. Cells shown are limited to those author-annotated as “bronchial goblet cell” or “mucus-secreting cell”. It can be seen that some cells secrete both MUC5AC and MUC5B to significant levels (e.g. clustered around 45 degree line on scatterplot), while others secrete one of MUC5B or MUC%A but little of the other (those on the axes). Sourced from the Human Lung Atlas database, found at <https://data.humancellatlas.org/hca-bio-networks/lung/atlases/lung-v1-0>; described in H.B. Schiller, et al. The Human Lung Cell Atlas: A High-Resolution Reference Map of the Human Lung in Health and Disease, Am J Respir Cell Mol Biol 61(1) (2019) 31-41.

| **NHBE donor (our reference)** | **Lot** | **Tissue Acquisition Number** | **Age** | **Sex** | **Race** |
| --- | --- | --- | --- | --- | --- |
| 1 | 18TL215671 | 36311 | 47 | F | H |
| 2 | 18TL127524 | 35773 | 69 | M | C |
| 3 | 0000646466 | 33261 | 38 | M | C |
| 4 | 404Z008 | 29910 | 65 | F | C |
| Summary |  |  | Mean age 55 | Ratio 50% M:F |  |
| **CHBE donor (our reference)** | **Lot** | **Tissue Acquisition Number** | **Age** | **Sex** | **Race** |
| 1 | 18TL179343 | 36100 | 69 | M | C |
| 2 | 18TL204559 | 36221 | 54 | F | B |
| 3 | 0000440551 | 28154 | 63 | M | B |
| 4 | 0000407341 | 27425 | 76 | F | C |

***Supplementary Table S1. List of donors of cells.***

| **Reagent** | **Volume (µl)** | **Final concentration** |
| --- | --- | --- |
| Dulbecco's Modified Eagle Medium | 500 | - |
| L-glutamine | 5 | - |
| Airway epithelial basal medium | 500 | - |
| Bovine Pituitary Extract | 2ml | 0.004 ml/ml |
| Epidermal Growth Factor (recombinant human) | 500 | 10 ng/ ml |
| Insulin (recombinant human) | 500 | 5 µg/ml |
| Hydrocortisone | 500 | 0.5 µg/ml |
| Epinephrine | 500 | 0.5 µg/ml |
| Transferrin (recombinant human) | 500 | 10 µg/ml |

**Additionally,** Retinoic acid (final concentration at 1:5000, *Sigma Aldrich, Dorset, UK*) was added just before use.

**Supplementary Table S2. Composition of bronchosphere culture media.**

**Supplementary Video S1 3D reconstruction showing mucus secretion in human bronchial organoids (included as separate .avi file).** (Photostiffened gel, 1 minute photocuring, 72 h incubation after photocuring.) 3D reconstruction from confocal (immuno)fluorescence image: of MUC5AC (Yellow) and MUC5AC (pink) along with cell nuclei (DAPI, blue).

| ***Antibody/stain name*** | ***Final Concentration*** | ***Manufacturer*** | ***Cat. No.*** |
| --- | --- | --- | --- |
| **Primary antibodies** | | | |
| Muc5ac | 1:100 | ThermoFisher, Paisley, UK | MA5-12178 |
| Muc5b | 1:200 | Elabscience, Texas, USA | EAB-15988-ELA |
| **Secondary antibodies** | | | |
| Goat anti-mouse AF488 | 1:500 | ThermoFisher, Paisley, UK | A21121 |
| Goat anti-mouse AF568 | 1:500 | ThermoFisher, Paisley, UK | A21144 |
| Goat anti-rabbit AF568 | 1:500 | ThermoFisher, Paisley, UK | A11036 |
| Goat anti-rabbit AF647 | 1:500 | ThermoFisher, Paisley, UK | A27040 |
| **Non-antibody stain** | | | |
| DAPI (4′,6-diamidino-2-phenylindole) | 1:1000 | ThermoFisher, Paisley, UK | D1306 |

**Suppl. Table S3. Antibodies and stains used in confocal imaging.**
